# Methodological determinants of signal quality in electrobulbogram recordings

**DOI:** 10.1101/2025.07.01.662502

**Authors:** Frans Nordén, Irene Zanettin, Tora Olsson, Artin Arshamian, Mikael Lundqvist, Fahimeh Darki, Johan N. Lundström

**Author notes:** Corresponding authors* Frans Nordén, MSc., Dept. of Clinical Neuroscience, Karolinska Institutet, Nobels väg 9, 17177 Stockholm, Sweden, Johan Lundström, PhD, Dept. of Clinical Neuroscience, Karolinska Institutet, Nobels väg 9, 17177 Stockholm, Sweden. Indicates shared senior author (Fahimeh Darki, Johan N. Lundström).

## Abstract

The electrobulbogram (EBG) is a new, non-invasive method for measuring the functional activity of the human olfactory bulb (OB). To date, the EBG has been used to assess how the OB process odor identity, valence, intensity, and it has shown promise as an early biomarker for Parkinson’s disease. However, current implementation of the EBG method depends on several methodological components, including subject specific co-registration of electrode positions through neuronavigation and EEG source reconstruction, which may limit accessibility for many research groups. In this study, we test the quality and reliability of the OB signal under different configurations to potentially remedy this. Specifically, we compare six EBG setups that vary in the use of subject-specific T1 scans versus a template head model, co-registered versus template electrode positions, and individualized versus template-based OB location. Our results indicate that strongest EBG signals are obtained when using subject-specific T1 scans in combination with co-registered electrode positions. However, we obtained significant EBG activity even when using a fully template-based configuration. Our anatomical analysis of OB location of 941 individuals reveals that in 86% of cases, the OB is centered within the spatial resolution bounds of the EEG source dipole, supporting the feasibility of detecting olfactory bulb signals without precise individual anatomical mapping using template coordinates. These findings suggest that while subject-specific configurations enhance signal quality, the EBG method remains robust enough to yield meaningful results even with less complex setups. This enables a broader adoption of the EBG method in both clinical and research settings.

## Introduction

The olfactory bulb (OB) is the first structure in the central olfactory system and plays a critical role in the initial processing of odor information. Although promising recent attempts of assessing OB activity using traditional neuroimaging methods have been made, assessing neural processing within the OB has remained challenging due to its location close to the sinuses. (Miao et al., 2021) We recently presented a new method that allows recording of valid and reliable neural signals from the human OB in a non-invasive fashion, the electrobulbogram (EBG) method (Iravani et al., 2020). The EBG is the only existing method to assess signal from the human OB with high temporal precision, but it is methodologically challenging to use due to its reliance on multiple costly techniques. Streamlining the EBG protocol to reduce both cost and the need for advanced technological expertise would enable broader use in experimental and clinical research.

The EBG method captures neural responses from the OB by applying source reconstruction techniques on recorded EEG signals from electrodes placed at the nasal bridge during odor delivery. Critically, the method traditionally requires knowledge of the anatomy and location of the OB. Thus, subject-specific electrode positions are co-registered with a neuronavigation system using individual T1 from MRI scans. The setup is then combined with an olfactometer that can deliver odor in synchrony with inhalation. The EBG method has been demonstrated to produce valid and reliable results (Iravani et al., 2020), to replicate well-established response patterns observed in non-human animal models (Iravani, et al., 2021a), and to distinguish between Parkinson patients and healthy age/gender-matched control individuals (Iravani, et al., 2021b). While the method has the potential to serve as a valuable tool in both research and clinical applications, its adoption has been limited; mainly because few laboratories possess all the necessary tools and infrastructure to acquire EBG according to the established protocol, restricting its availability, particularly in clinical settings.

To increase availability of the EBG method, we systematically assess each major component’s contribution to the overall sensitivity of the measure. Specifically, we provide guidance on whether to use participant-specific T1 MR images or whether a population average template T1 would suffice, whether co-registered electrode coordinates obtained from a neuronavigation system is needed or whether template electrode positions would suffice, and finally, how the contribution of individually mapped OB coordinates compares to standardized OB source coordinates, obtained from group average data.

## Method

### EBG evaluation

#### Participants

A total of 53 individuals, self-described as healthy with a normal sense of smell, were initially recruited for the comparative analysis. Following the exclusion of seven participants due to reasons outlined in the preprocessing section, the final dataset consisted of 46 participants (mean age: 30 years, SD ±8.8; 27 women). To confirm that all participants had a functional sense of smell, an odor screening test was conducted in which they were asked to identify five different odors using a four-alternative odor identification task (Iravani et al., 2020; Iravani et al., 2021c). A minimum of three correct responses was required to ensure the absence of functional anosmia or severe hyposmia. The study was approved by the Swedish National Ethical Permission Board (Etikprövningsnämnden, EPN: 2017/2332-31/1) and adhered to the principles of the Declaration of Helsinki. Before participating, all individuals provided written informed consent regarding the study’s purpose and procedures.

#### Testing procedure

The experiment was conducted in a sound-attenuated and well-ventilated recording booth, specifically designed for odor testing, ensuring a controlled environment with minimal background scents. To eliminate potential auditory cues from the olfactometer and odor delivery system, participants wore earplugs and headphones that played low-volume white noise throughout the session. PsychoPy 3 (Peirce et al., 2019) was used to control event timing and stimulus presentation.

Delivery of odor stimuli was aligned with individual nasal inspiration. Participants were therefore instructed to breathe naturally through their nose throughout the experiment. The experiment comprised four blocks, each lasting 15 minutes, with short breaks in-between. Each block contained 35 trials, resulting in a total of 140 experimental trials per participant. These trials included three different odors, each presented at two concentrations, and a Clean air condition. Odor presentation was randomized across blocks to prevent predictability.

Following each trial, participants rated the intensity and pleasantness of the stimulus using a 0 to 100 scale, where 0 indicated no perception or very unpleasant, and 100 represented very intense or very pleasant (these ratings are not assessed in this manuscript). To minimize odor habituation, the inter-trial interval (ITI) was jittered but maintained at a minimum of 14 seconds.

#### Odor delivery and odor stimuli

Three odorants were used: n-Butanol (an alcohol, Fisher Chemicals, CAS 71-36-3), 5-Nonanone (a ketone, Sigma-Aldrich, CAS 502-56-7), and Undecanal (an aldehyde, Sigma-Aldrich, CAS 112-44-7). Each odorant was dissolved in diethyl phthalate (99.5% pure, Sigma-Aldrich, CAS 84-66-2) and prepared at two concentration levels. The low-concentration solutions were diluted to 0.8%, 1.3%, and 1.4%, while the high-concentration solutions were prepared at 4%, 18%, and 17%, respectively. We used different concentrations of the odorants because the experiment’s main aim was to assess how the low-level olfactory system process odor intensity; these experimental data are repurposed to assess our methodological question. Results of the main experimental aims are presented elsewhere (Nordén et al., 2025). A no-odor condition was also included, where trials consisted solely of clean air replacing the ongoing airflow.

Odor presentation was controlled using a computer-controlled olfactometer (Lundström et al., 2010), delivering stimuli birhinally for two seconds per trial in a sniff-triggered design. A thermopod sensor (sampling rate: 400 Hz; PowerLab 16/35, ADInstruments, Colorado) was placed inside the nostrils to monitor breathing patterns based on intranasal temperature. The recorded breathing signal was used to establish an individualized threshold, where the participant’s breathing cycle, unbeknown to them, triggering the olfactometer precisely at the onset of inhalation. This synchronization between odor delivery and nasal inspiration was applied to minimize attention-related EEG artifacts. An overview of a trial can be seen in Figure 1.

**Figure 1.**
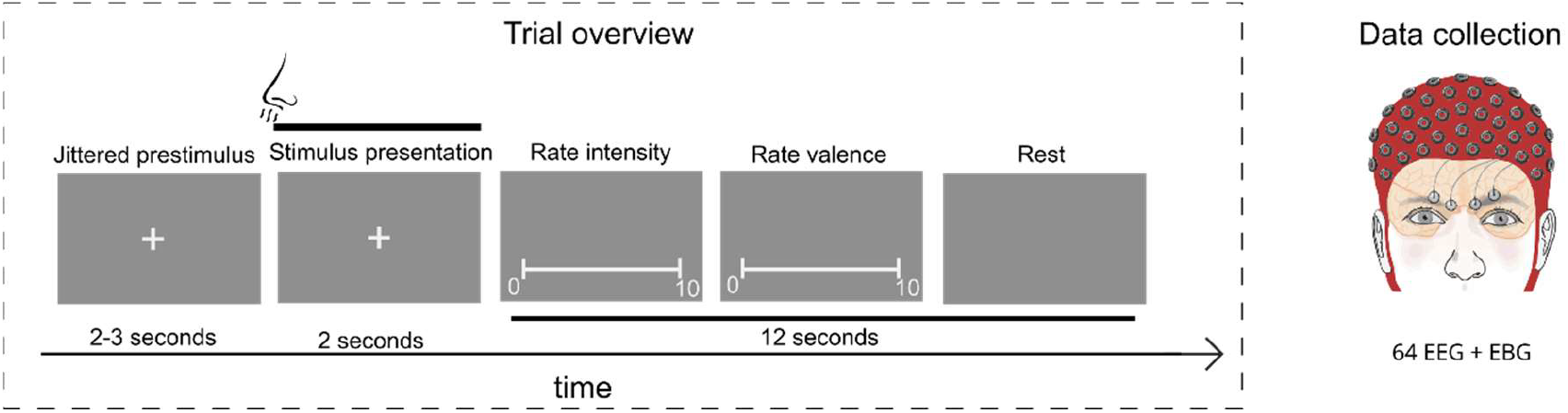
Overview of the stimulus presentation and the location of the EBG electrodes.

To prevent participants from detecting any tactile airflow changes at odor onset, a birhinal background airflow of 0.3 liters per minute of clean air was continuously maintained, with either odorized airflows or clean air of 2.7 liters per minute. The olfactometer introduced a latency of approximately 150 milliseconds, accounting for the trigger-time and time required for the odor to travel through the tubing and reach the nasal epithelium. Prior to the experiment, this delay was verified using a photo-ionization detector (Aurora Scientific, Ontario), and all subsequent analyses were adjusted accordingly. As a result, time zero in all reported data reflects the moment the odor entered the nostrils.

#### EEG and EBG recordings

Neural activity was continuously recorded at a sampling rate of 512 Hz using a setup consisting of 64 electroencephalography (EEG) electrodes and 4 electrobulbogram (EBG) electrodes (ActiveTwo, BioSemi, Amsterdam, The Netherlands). EEG electrodes were positioned in accordance with the international 10/20 system while EBG electrodes were placed just above the eyebrows to capture olfactory-related neural activity. The precise spatial coordinates of each electrode were digitized in stereotactic space using a neuronavigation system (BrainSight, Rogue Research, Montreal, Canada). This process involved registering fiducial landmarks, including the nasion, left and right preauricular points, as well as the central locations of each EEG and EBG electrode to ensure accurate alignment with the individual’s head anatomy.

Throughout the recording session, real-time signal processing was applied, with the neural data high-pass filtered at 0.10 Hz and low-pass filtered at 100 Hz using ActiView software (BioSemi, Amsterdam, The Netherlands). Before data acquisition, electrode impedance was assessed and any electrode exhibiting an offset greater than 40 mV was adjusted to fall within the acceptable threshold, ensuring optimal signal quality.

#### Preprocessing

Before preprocessing, five participants were excluded due to minimal variability in their ratings, as they consistently rated all odors as nearly identical, indicating a dysfunction or malingering. This left 48 participants for further analysis.

For these remaining subjects, the neural data were segmented into 5000 ms epochs, spanning 1000 ms before stimulus onset to 4000 ms post-stimulus, and subsequently re-referenced to the average activity of all electrodes. A high-pass filter at 1 Hz was applied to remove slow drifts, while a 50 Hz notch filter was used to eliminate power line interference. Faulty electrodes were identified through visual inspection and corrected using spline interpolation.

To address artifacts, trials with excessive muscle activity were removed by extracting z-scored Hilbert transform amplitude values and applying a threshold of 7. Additionally, eye blinks were detected and removed using Independent Component Analysis (ICA) with the InfoMax algorithm. Two participants were excluded at this stage due to insufficient remaining trials, having lost more than half of their data during artifact rejection. Following preprocessing, the final dataset consisted of 46 participants, with an average of 123 ± 6.84 trials available for statistical analysis.

#### Source reconstruction

We wanted to evaluate how six different methodological configurations affect the results of the EBG method. The EBG method (Ground truth) use subject-specific T1 scans, electrode positions co-registered with the individual head model, and OB location chosen from the subject specific T1 and T2 scans. For source reconstruction, structural MRIs are segmented into five aspects, namely CSF, gray matter, white matter, scalp, and skull with conductance’s [0.43, 0.01, 1.79, 0.33, 0.14] (Hallez et al., 2007) and the forward problem is solved using the finite element method implemented in Simbio (Vorwerk et al., 2018). A cortex model built on icosahedrons with resolution 7 based on individual structural MRI is then generated in Freesurfer and used as the source model. This source model is subsequently transformed into MNI stereotactic space to ensure that identical regions are used for all participants. This source model is then used to attain the source activity through solving the inverse problem with eLORETA. The regularization parameter is set to 10% and singular value decomposition is used to project the source activity along the principal axis.

The other configurations assessed here were reductions from the ground truth. The second configuration was the same as the ground truth but with the OB locations linearly transformed from the average OB location found in the second part of the analysis (x +/-3, y+42, z-31). The third and fourth configurations correspond to the first and second configurations but with electrode positions taken from the BioSemi 64-10/20 template with the four EBG electrodes added with locations chosen above the eyebrows on the MNI152 T1 template.

The fifth and sixth configurations utilized a methodology based on (Fuchs et al., 2007) and utilized in (Nordén et al., 2024) where a template T1 scan was used to solve the forward problem. A cortex-based head model was generated in the same way as for the other configurations, but instead of subject specific T1 scans it was based on the MNI152 template. The difference between the fifth and sixth configuration is that for the fifth configuration the electrode positions were co-registered to the participants head and then fitted manually to the template head model and for the sixth configuration the template electrode layout with the four added EBG-electrodes were used. The location of the OB was chosen based on the MRI based analysis and the source reconstruction was performed with Fieldtrip toolbox 2023 (Oostenveld et al., 2011) within MATLAB 2023a .

#### Evaluation

To determine which of the configurations demonstrated the best performance, we benchmarked them against activity in the early gamma band (100-300 ms after the odor enters the nose). This neural response is viewed as the hallmark OB response and was first observed in intracranial recordings of the OB (Hughes et al., 1969) and later replicated using the non-invasive EBG method (Iravani et al., 2020).

We selected three different measures to evaluate the results. The first measure focused on the signal-to-noise ratio (SNR), comparing the neural response to Odor stimuli versus Clean air in the early gamma band. The second measure evaluated statistical significance, expressed as a *p*-value, derived from the contrast between odor and Clean air conditions in the early gamma band. This analysis uses cluster-based permutation test with 5,000 permutations, and the results were cluster-corrected using the weighted cluster mass algorithm (Hayasaka & Nichols, 2004). Lastly, the third measure examined the effect size within the same region where the significance testing was performed, providing insight into the magnitude of the observed differences.

#### Probability map of the OB location

The ability of standard group-derived OB coordinates to capture true OB signal is tightly coupled with the underlying variability of the anatomical location of the human OBs; a measure not previously assessed. To define the typical spatial variability of the OB in an adult healthy population, we aimed to generate a probabilistic map of the OB’s location using data from the Human Connectome Project Young Adult (HCP-YA). The HCP-YA dataset includes high-resolution T1- and T2-weighted MRI image data from a cohort of ∼1100 healthy individuals aged 22 to 35 years).(Van Essen et al., 2013)

The OB was segmented and identified in each individual using the automatic segmentation tool (Desser et al., 2021), which employs a 3D U-Net convolutional neural network. This model performs automated OB segmentation based on the combined T1- and T2-weighted images; an approach shown to achieve accuracy comparable to manual segmentations by expert raters. For details about the segmentation tool pipeline see (Desser et al., 2021). All preprocessing, segmentation, and postprocessing steps were performed using Python version 3.7.

Automated OB segmentation was performed on 1101 subjects with both T1- and T2-weighted images in native subject space. To ensure the anatomical plausibility and reliability of OB segmentations, we implemented a rigorous postprocessing quality control procedure to identify and exclude erroneous results. First, participants were excluded if the voxel distance (*D)* between the left and right segmentations was either *D* ≤ 0 or *D* ≥ 8, indicating no separation or unrealistically large separation between the left and right bulbs. Second, participants were discarded if both OBs were located within the same hemisphere. Third, participants were excluded if one bulb had volume <= 15 mm^3^ and the absolute volume difference between the left and right OB volume exceeded 10 mm^3^, suggesting segmentation failure on one side.

Applying these criteria resulted in the exclusion of 160 participants, leaving a total of 941 segmented OBs in the final dataset. To construct a probabilistic map reflecting the typical spatial distribution of the OB across healthy individuals, the OB masks in MNI space were combined across all 941 subjects, and the resulting sum was divided by the total number of participants. Each voxel of the resulting probabilistic map indicates the proportion of participants in whom the voxel was classified as part of the OB.

## Results

### Olfactory bulb location is consistent across individuals

To test the critical components of the electrobulbogram (EBG) method, we first evaluated the location of the olfactory bulb in a larger population. For this part, we used T1- and T2-weighted MRI scans from the young adult dataset of the Human Connectome Project (HCP). The region where the majority of subjects’ OBs were found is visualized in Figure 2. This region corresponded to center coordinates in the MNI space of the left OB at (x - 3, y + 42, z - 31) and the right OB at (x + 3, y + 42, z – 31). A total of 68% of the tested subject’s OB showed an anatomical overlap with these coordinates, indicating a high degree of spatial consistency across individuals in OB location.

**Figure 2.**
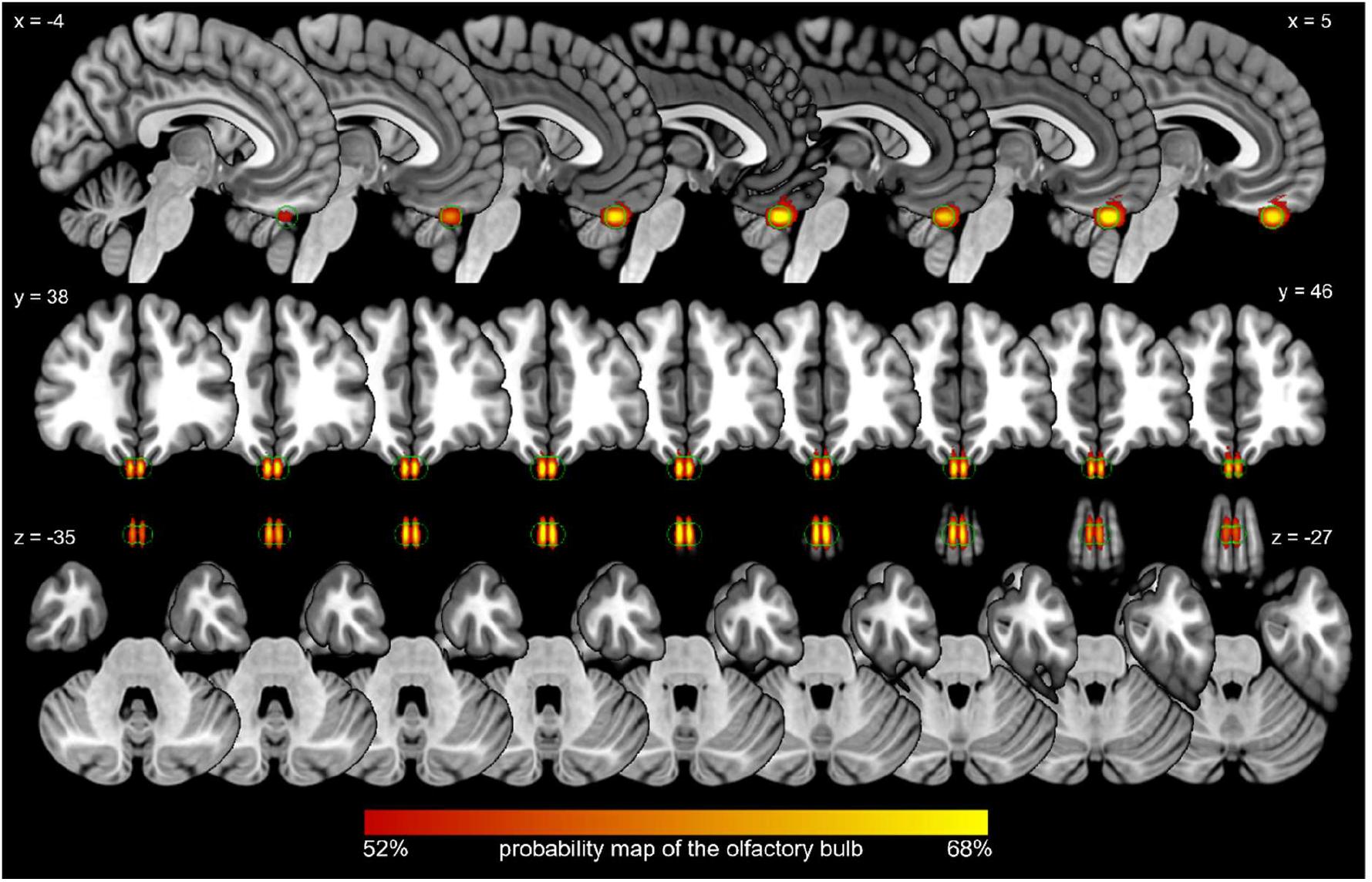
Probabilistic map of olfactory bulb location. Based on the automatic segmentation of left and right olfactory bulbs of 941 healthy individuals from the human connectome project database. Colormap indicates the proportion of participants in whom the voxels were classified as part of the olfactory bulb.

Further examination in 3D space revealed that the location of the OB was relatively consistent across subjects (Figure 3) in most axis. Specifically, along the x-axis, the OBs exhibited minimal lateral deviation, with two standard deviations encompassing a radius of only 2 mm from the mean OB location. Along the y-axis, the variability was somewhat greater, with a radius of 6 mm, indicating slightly more positional variation in the anterior-posterior direction. The most substantial variation was observed along the z-axis where the OB locations extended within a 9 mm radius from the average position. Despite this variation, the majority of subjects’ OBs were located within a relatively confined spatial region, reinforcing the reliability of using an average OB position in our source model approximation. More specifically, when considering the EEG dipole radius, in our case 12mm in diameter, approximately 86% percent of these 941 participants had an OB within the sphere when using the average coordinates (left OB 86.32%, right OB 86.56%).

**Figure 3.**
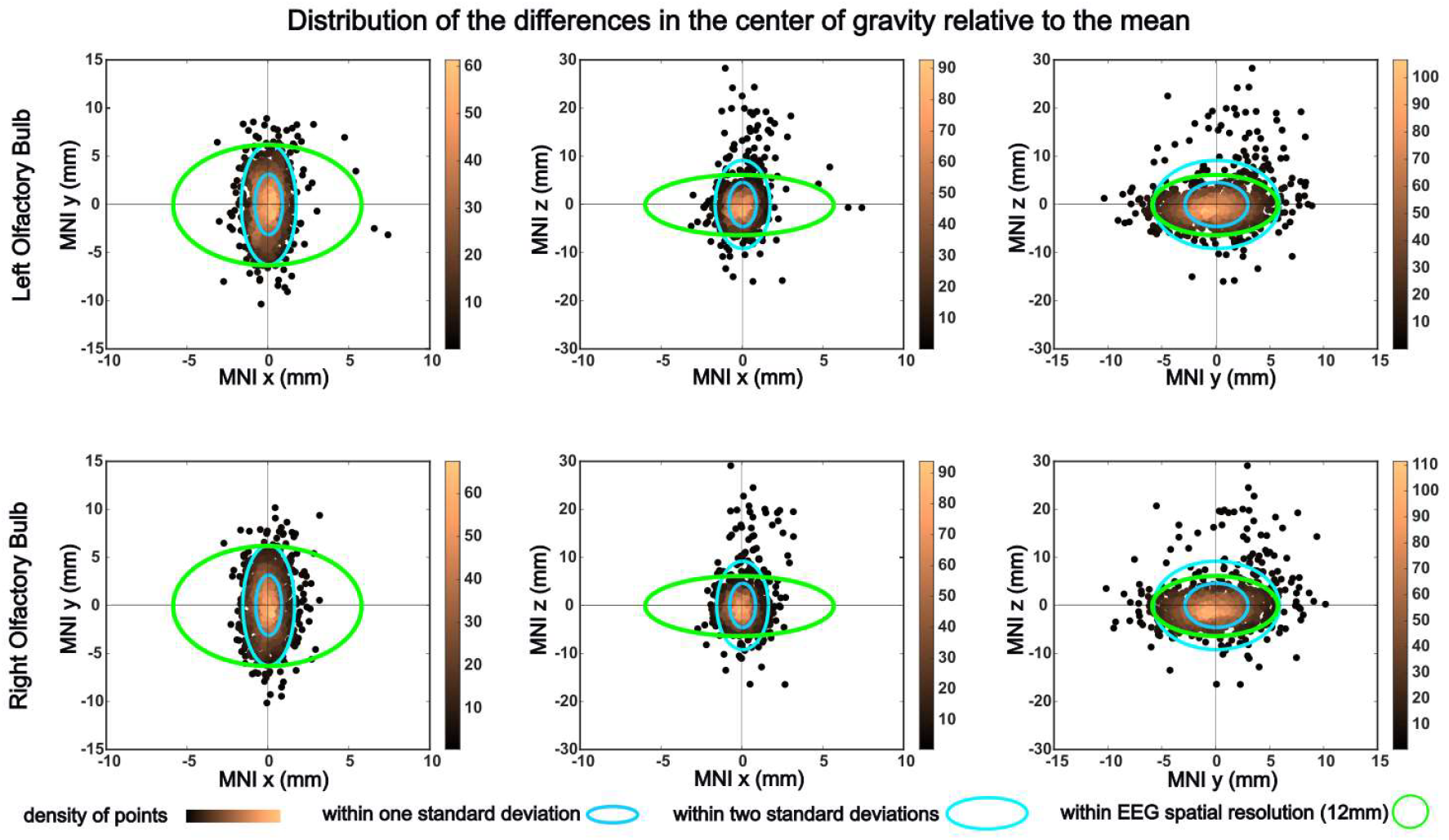
Typical spatial variability of the olfactory bulb location. Individual variation in olfactory bulb location reveals that a difference of 1–2 cm in olfactory bulb center position falls within two standard deviations of the observed distribution.

### EBG signal is detected in all configurations

To further determine the necessary configurations for reliable electrobulbogram (EBG) activity measurement, we tested six different configurations, each varying in their use of subject-specific or template-based anatomical data, co-registered electrodes through neuronavigation, and OB localization methods. The following configurations were tested:

1. Ground truth: subject specific T1, neuronavigation, and manually extracted OB location (T1, NN, loc)
2. Same as 1, but with OB location based on the average location in MNI coordinates (T1, NN, -loc)
3. Subject specific T1, no neuronavigation, manually extracted OB location (T1, -NN, loc)
4. Same as 3 but with OB location as in 2 (T1, -NN, -loc)
5. MNI template T1 and neuronavigation (-T1, NN, -loc)
6. MNI template T1, no neuronavigation, average OB location (-T1, -NN, -loc)

Evaluating the signal-to-noise ratio (SNR) in the selected region, we observe that it is considerably higher in the configurations that incorporate T1-weighted MRI scans and co-registered electrode positions (Supplementary Table S2). While all methods produce SNR values above zero, the difference between this and the other configurations was striking, suggesting that co-registered electrode positions play a crucial role in enhancing signal clarity (Figure 4).

**Figure 4.**
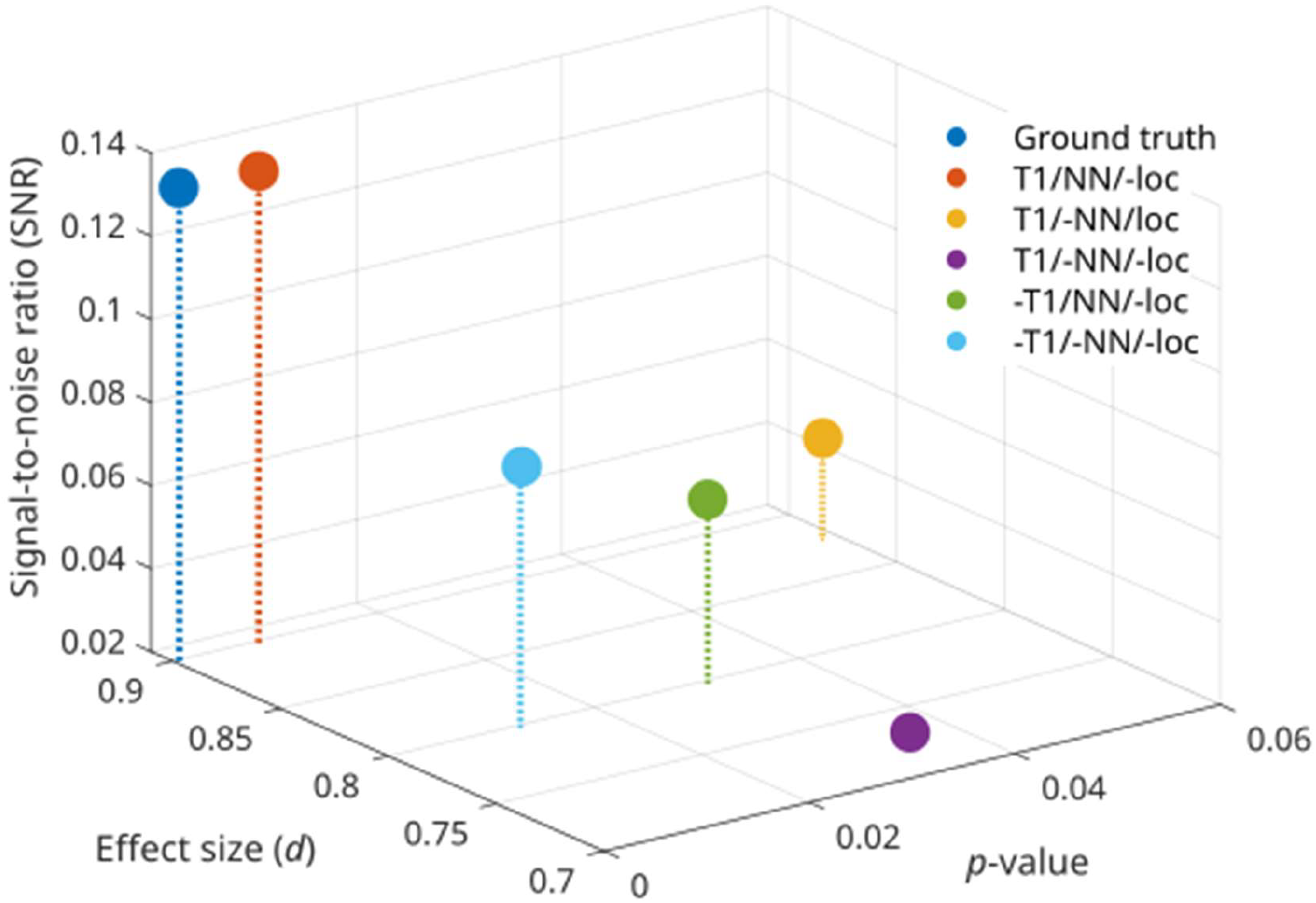
Evaluation of the six configurations in the early gamma region based on three criteria: Signal-to-noise ratio (SNR), effect size and p-value. Ground truth shows the strongest results in all three criteria closely followed by the configuration without manually chosen OB location. All configurations do, however, show positive SNR and strong effect size measured with Cohen’s D. The only configuration not reaching the statistical significance level when correcting for multiple comparisons was the one using subject specific T1 but not subject specific co-registered electrode positions.

When examining statistical significance through *p*-values, we find that all configurations, except one, achieve values below the standard significance threshold (*p* < .05). The exception is Configuration 3 (T1, -NN, loc), which yields a *p*-value of .06, slightly above the threshold but still close to statistical significance. The best-performing configurations, Configuration 1 (T1, NN, loc) and Configuration 2 (T1, NN, -loc), demonstrate statistically significant results with *p*- values of .003 and .004, while Configuration 4 (T1, -NN, -loc), which shares similarities with Configuration 3 (T1, -NN, loc), but relies on a template-based OB location, demonstrated lower values (*p* = .03). The two configurations that omit subject-specific T1 scans, Configuration 5 (- T1, NN, -loc) and 6 (-T1, -NN, -loc), perform similarly, with *p*-values that fall within the significance range (*p* = .03, *p* = .01).

The time frequency responses (Supplementary Figure S1) show that the responses differ between configurations. There is a clear difference between the configurations using a subject specific T1 and the ones that do not. In the ones with template T1, we see a stronger activation in lower gamma while we in the ones using a subject specific one has activation that is more spread-out in the frequency domain. Ground truth (T1, NN, loc) and Configuration 2 (T1, NN,

-loc) combines both the higher gamma activation with the lower and therefore achieves higher statistical significance than the other configurations.

Assessing effect sizes, we find that all configurations perform reasonably well, with values exceeding Cohen’s *d* = 0.7, indicating a strong effect across conditions. The highest effect sizes are observed in Configuration 1 and 2, both reaching 0.9, and Configuration 3, reaching 0.87, suggesting a particularly robust response in these configurations. In contrast, Methods 5 and 6 exhibit slightly lower effect sizes of 0.8, while Configuration 4 shows the weakest effect, with a value of 0.72.

## Discussion

Here, we aimed to answer two questions: First, how six different methodological choices affect the ability of the electrobulbogram (EBG) method to capture the neural signal from the OB; and second, how much individual variation exists in the location of the human olfactory bulb (OB). We provide evidence that 86% of subjects have an OB centered within the spatial resolution of the EEG source dipole. We further provide evidence for the importance of subject-specific anatomical data in combination with co-registered electrode positions in improving signal quality, statistical reliability, and effect size in EBG measurement. Our data also shows that a simple configuration using only a template-based head model and template electrode positions renders significant results in the early gamma region.

We found that accurate co-registration of EEG and EBG electrodes was one of the most critical factors in obtaining strong results when applying the EBG method. This became particularly evident when comparing the ground truth configuration to Configuration 3, which used the same setup, but without co-registered electrodes. While the ground truth configuration yielded highly significant results and a strong signal-to-noise ratio (SNR), Configuration 3 failed to reach statistical significance and exhibited a substantially lower SNR. This discrepancy is not surprising given that the EEG cap often needs to be positioned slightly further back to accommodate the EBG electrodes. Such an adjustment inevitably alters the placement of all electrodes on the individual level, introducing spatial misalignment that is not accounted for in a template-based electrode layout. Additionally, individual variations in head shape further contribute to misalignment, which is not captured when using standardized electrode templates.

One could argue that the spatial displacement of electrodes is smaller than the intrinsic spatial resolution of EEG, and while it might be true, it is also important to consider the small size of the OB. Unlike cortical sources, where minor shifts in electrode positioning may have negligible effects, the OB is a compact structure, making precise electrode placement especially crucial for detecting its activity. Even small deviations in electrode location could lead to signal attenuation, increased noise, or spatial smearing, ultimately diminishing the reliability of the recorded EBG signals. Our findings highlight the importance of individualized co-registration of electrode positions in ensuring accurate localization of OB activity and suggest that electrode placement errors could be a major limiting factor in the robustness of the EBG method.

Examining additional factors that influence the strength of the results, it is evident that the second most significant factor, closely tied to accurate electrode co-registration, is the use of a subject-specific T1 scan for head and source modeling. This is consistent with the current literature that concludes that there is a clear improvement with subject specific T1s (Depuydt et al., 2024). The two configurations that incorporated both subject-specific T1 scans and co-registered electrode locations consistently produced the strongest results across all our metrics.

It is worth noting that co-registered electrode locations did not yield noticeable improvement when using a template-based T1 configuration. This can likely be attributed to the inherent limitations of template-based head models, which tend to produce less precise and more spatially smeared source reconstruction (Depuydt et al., 2024). As a result, this smearing may act as a smoothing effect and reduce the influence of minor variations in electrode placement, effectively diminishing the benefits of precise co-registration.

While our findings highlight the advantages of individualized anatomical modeling, they also demonstrate that meaningful effects can still be detected using alternative configurations. This suggests that, although subject-specific head models and precise electrode placement enhance the robustness of the analysis, significant findings remain achievable even when relying on template-based approaches.

Given the importance of accurate electrode localization, it is surprising that manually identifying the OB does not have a stronger effect on the results. This is particularly unexpected considering that our initial analysis revealed that 14% of participants have an OB located outside the spatial resolution of EEG. In theory, losing the signal from such a proportion of subjects should reduce statistical significance, yet this effect is not clearly reflected in the comparison of *p*-values, SNR, or obtained effect size. This might be explained by head model based volume conduction that is not accounted for by the EEG spatial resolution (Srinivasan et al., 2007).

Our findings suggest that, regardless of the chosen configuration among those tested, the likelihood of detecting a signal from the OB remains high. While this likelihood naturally improves with methodologies designed for greater accuracy, certain factors, such as the use of subject-specific OB localization, do not appear to have a significant impact. If there is a trade-off to be made, sampling from more participants is likely of greater importance than maximizing signal quality.

By identifying key factors influencing EBG signal quality, we aim to make the EBG method more accessible to a broader range of laboratories, facilitating non-invasive recordings from the OB. This advancement paves the way for a deeper understanding of the neural mechanisms underlying human olfactory processing by providing clear methodological guidance. These findings hold promise for olfactory-related translational research and can be readily integrated into clinical settings to detect central olfactory processing changes associated with neurological and neurodegenerative diseases.

## Data availability

All anonymized data and scripts to reproduce the results are available at https://osf.io/3zk2g/?view_only=ac8a5462c0cd419fb203a941e9887d82

## Author contributions

Conceptualization: FN, ML, AA, FD, JNL Methodology: FN, IZ, ML, AA, FD, JNL Investigation: FN, IZ, TO

Visualization: FN, TO, FD Supervision: FD, JNL Writing—original draft: FN, TO, AA

Writing—review & editing: FN, IZ, TO, AA, ML, FD, JNL

