## Supplementary materials for "Methodological determinants of signal quality in electrobulbogram recordings"

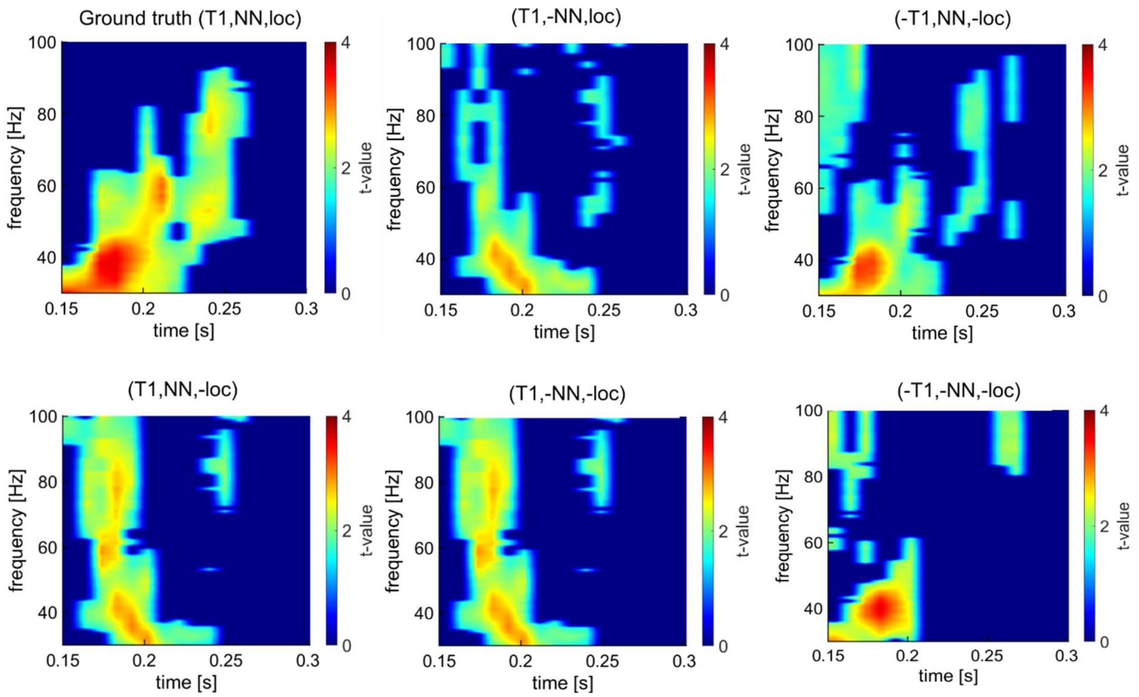


**Supplementary figure 1: Early gamma evoked response in the six different configurations. Ground truth provides both the biggest area of activity and the strongest intensity.**

**Supplementary table 1: Signal to noise ratio in the OB early gamma region.**

| SNR gamma | Participant T1 with manual OB position | Participant T1 with template OB position | Template headmodel |
|---|---|---|---|
| Neuronavigation | 0.13 | 0.14 | 0.067 |
| Template electrodes | 0.047 | 0.024 | 0.082 |
